# Origin of biogeographically distinct ecotypes during laboratory evolution

**DOI:** 10.1101/2023.05.19.541524

**Authors:** Jacob J. Valenzuela, Selva Rupa Christinal Immanuel, James Wilson, Serdar Turkarslan, Maryann Ruiz, Sean M. Gibbons, Kristopher A. Hunt, Manfred Auer, Marcin Zemla, David A. Stahl, Nitin S. Baliga

## Abstract

Resource partitioning within microbial communities is central to their incredible productivity, including over 1 gigaton of annual methane emissions through syntrophic interactions^1^. Here, we show how isogenic strains of a sulfate reducing bacterium (*Desulfovibrio vulgaris*, Dv) and a methanogen (*Methanococcus maripaludis*, Mm) underwent evolutionary diversification over 300-1,000 generations in a purely planktonic environmental^2–4^ context giving rise to coexisting ecotypes that could partition resources and improve overall stability, cooperativity, and productivity in a simulated subsurface environment. We discovered that mutations in just 15 Dv and 7 Mm genes gave rise to ecotypes within each species that were spatially enriched between sediment and planktonic phases over the course of only a few generations after transferring the evolved populations to a fluidized bed reactor (FBR). While lactate utilization by Dv in the attached community was significantly greater, the resulting H_2_ was partially consumed by low affinity hydrogenases in Mm within the same attached phase. The unutilized H_2_ was scavenged by high affinity hydrogenases in the planktonic phase Mm, generating copious amounts of methane and higher ratio of Mm to Dv. Our findings show how a handful of mutations that arise in one environmental context can drive resource partitioning by ecologically differentiated variants in another environmental context, whose interplay synergistically improves productivity of the entire mutualistic community.

## INTRODUCTION

Microbial communities drive all biogeochemical processes on Earth through spatiotemporal resource partitioning among co-existing members or ecotypes that are ecologically differentiated by their preference for a particular environmental context like seasonal fluctuations or physical attributes like free-floating (planktonic) or attached lifestyles^5,6^. Ecotype differentiation, which leads to resource partitioning, has long been known to be a key mechanism by which microbes “divide and conquer” diverse niches. There are numerous examples of how even within a species (e.g., species of *Prochlorococcus*^7^, *Vibrio*^6^, etc.) resources are partitioned in a single habitat through ecotype differentiation. Each ecotype is specialized to maximally utilize some subset of resources, each occupying a different niche. In fact, only recently using comparative genomics have we gained a better understanding of how finely niche space is divided, even in what might appear to be a homogeneous environment. This adaptive diversification is the underpinnings of the remarkable diversity of microbial life now being more revealed through the developing science of metagenomics. It has recently been estimated that 20-80% of the cells in subsurface environments exist as biofilms exhibiting, relative to their planktonic counterparts, distinct physiologies and phenotypes, such as increased resistance to antimicrobials, heavy metals, desiccation and substrate deprivation^8–11^. The local genotype interactions among these ecologically differentiated strains confer emergent properties such as stability and cooperativity to the microbial community^12^. While there is evidence that extended periods of evolution may eventually drive sympatric speciation of ecologically differentiated strains^13–15^, there is scant evidence of early events, mechanisms, and time frames over which ecological differentiation manifests in a microbial community of clonal isolates of two or more organisms.

We have discovered early events in ecological differentiation of a nascent community of sulfate reducing bacteria (**SRB**) and methanogens engaged in a type of mutualism called syntrophy, which is responsible for transformation of >1 gigaton/yr of C into CH_4_^1^. This particular type of syntrophy can be either obligate or facultative, depending on the availability of sulfate. Sulfate respiration (**SR**) is energetically favorable compared to syntrophic growth, which requires interaction with a methanogen in a sulfate-depleted and an energy limited anaerobic ecological niche^16–21^. The two-organism synthetic community (**SynCom**) of a SRB, *Desulfovibrio vulgaris* Hildenborough (**Dv**)^22^, and an archaeal methanogen, *Methanococcus maripaludis* S2 (**Mm**)^23^, has been used as a model to study syntrophic interactions in the laboratory^2–4,19–21,24,25^. When co-cultured in the absence of sulfate, Dv oxidizes lactate to acetate, formate and H_2_, but the build-up of H_2_ partial pressure eventually becomes growth inhibitory. Consumption of H_2_ or formate by Mm to produce methane (i.e., hydrogenotrophic methanogenesis) supports its growth, while reducing H_2_ partial pressure and alleviating Dv growth inhibition, making the overall reaction energetically favorable^16,26^. In so doing, together the SRB and methanogen are able to access a niche previously uninhabitable to both independently, and together drive one of the major processes of methane production on Earth.

Early on in the laboratory evolution of syntrophic interactions between Dv and Mm there were periods of instability and extinction events, but stability emerged within 300 generations significantly improving growth rate and yield^3^. Subsequent analysis discovered that while each evolved partner had individually contributed to improvements in growth characteristics, pairings of evolved clonal isolates of Dv and Mm from the 1,000^th^ generation synergistically improved overall growth characteristics. However, maximal improvements in growth rate and yield were observed in minimal assemblages of the syntrophic community that were obtained through end-point dilution of the evolved community from the 1,000^th^ generation, suggesting that overall growth improvements had emerged from the interplay of multiple variants across the two organisms, including low frequency variants that had retained SR capability^2,4^.

Using custom fluidized bed reactors (**FBRs**) to simulate an ecologically relevant subsurface environmental context for syntrophic interactions in soil (i.e., attached to sediment and free-floating in groundwater) we have investigated how the interplay between variants of vastly different abundance and physiological capabilities had contributed to improved growth characteristics of the minimal assemblage of the laboratory evolved community. Specifically, we asked whether diversification of genotypes in one environmental context (i.e., planktonic growth) could give rise to genotypes that might thrive in a separate environmental context^27,28^ (i.e., attached to sediment) to improve overall community characteristics? We have performed longitudinal assessments of changes in genetic diversity and transcriptome responses of Dv and Mm across attached and planktonic phases in the context of overall growth and productivity of the SynCom (i.e., lactate utilization and methane production). Further, by developing the first genome scale metabolic network models for an attached and planktonic syntrophic communities, we investigated dynamic changes in metabolic flux states across the planktonic and sediment phases. In so doing, we have followed the segregation of variants that emerged at the origin of a nascent microbial mutualism into biogeographically interdependent sub-communities, quantifying both the emergence and maintenance of new ecotypes. We have observed for the first time how mutations in a few genes across the two organisms mechanistically led to ecotype differentiation, driving resource partitioning and improving overall productivity of the mutualistic microbial community.

## METHODS

### FBR setup, operation, and inoculation with SynCom

Anaerobic FBRs were arrayed in triplicate in a temperature-controlled room (∼30°C) and filled with 75 g of 210 - 297 μm crushed quartz (Sigma-Aldrich: 50-70 mesh Quartz) to act as the sediment bed for microbial attachment. Reactors were filled with sterile lactate media and peristaltic pumps were used for recirculation to generate fluidization at 350 mL ⋅ min^-1^ with a duty cycle of 1 hour on per 2 hour period. Fresh lactate media was added (0.13 mL ⋅ min^-1^) to each reactor at the same duty cycle as the fluidization pumps. A simplified SynCom of Dv and Mm (EPD9) was sourced from a previous study ^2–4^ in which it was obtained through end-point dilution of a laboratory evolved co-culture after 1,000 generations of obligate syntrophic growth. A glycerol stock of EPD9 was revived in anaerobic Balch tubes containing lactate media and subjected to four successive transfers to reestablish growth prior to inoculation of each FBR. FBRs were operated in batch mode for approximately 48 hours to allow biomass accumulation before switching to fluidization mode for the rest of the experiment.

### Sample collection and analysis

An aliquot from the liquid phase of each FBR (“planktonic samples”) were used to measure optical density as a proxy for growth, lactate, acetate, pH, dissolved oxygen, and harvest cell pellets for DNA, RNA, and protein extractions (**Supplemental Table 1**). Lactate and acetate levels were quantified with ion chromatography as described previously^29^. Sediment samples were collected anaerobically on ice, washed twice with 10 mL of pre-chilled (4°C) PBS, split into aliquots, and stored at -20°C for DNA, RNA, and protein extractions. FBR headspace was sampled and stored in anaerobic vials containing 100% N_2_ for quantification of methane using gas chromatography.

### DNA, RNA, and protein extractions

Nucleic acid was extracted from planktonic and sediment samples using the MasterPure Complete DNA and RNA Purification Kit. DNA samples were treated with RNase and RNA samples were treated with DNase. All samples were quantified using the Qubit DNA or RNA High Sensitivity (HS) kits (Invitrogen) prior to sequencing. Protein extractions were performed according to Thermo Scientific B-PER Complete Bacterial Protein Extraction Reagent protocol. After protein was extracted from the attached community, the remaining sediment was dried and weighed for mass normalization.

### DNA and RNA library preparation and sequencing

DNA sequencing libraries were prepared using the Nextera XT DNA library prep kit, while RNA samples were rRNA depleted using RiboZero Plus kit followed by library construction using the Truseq Stranded Total RNA library prep kit. Both DNA and RNA libraries were sequenced using a mid-output 2 × 75 bp kit for 150 bp fragments using the Illumina NextSeq 500 platform.

### Identification of mutations

Mutations were determined using a custom sequence alignment and variant calling pipeline described previously^4^ (see https://github.com/sturkarslan/evolution-of-syntrophy and supplementary methods).

### Scanning electron microscopy

Morphology of Dv-*galU*_*P32S*_ mutant from a 300^th^ generation of an evolved co-culture^2–4^ was visualized with Hitachi S5000 Scanning Electron Microscope (Hitachi High Technologies America Inc, CA, USA).

### Gene expression analysis

Paired-end Illumina reads were processed using TrimGalore version 0.4.3_30_ and aligned to concatenated reference genomes of *Desulfovibrio vulgaris* Hildenborough (ASM19575v1) and *Methanococcus maripaludis* strain S2 (ASM1158v1) Transcript abundance was quantified using kallisto v0.44.0^31^ followed by separation of Dv and Mm reads prior to differential gene expression analysis using DESeq2 package v1.22.2^32^ in R.

### Metabolic model refinement and integration

Genome-scale metabolic models of Dv (iJF744)^33^ and Mm (iMR539)^34^ were updated and integrated into a single syntrophy model (iSI1283) by interlinking through known metabolite exchanges^19^. All metabolic model files are available for download from our GitHub repository https://github.com/baliga-lab/SynCom-model-for-DvH-and-Mmp.

### Flux predictions using state-specific models and analysis

iSI1283 was contextualized using planktonic and sediment attached phase gene expression data for all days using the GIMME algorithm^35^. We applied the constraint-based method for simulating the metabolic steady-state of iSI1283 using flux-balance analysis (FBA)^35,36^. The initial validation steps involved checking the capacity of the model to produce biomass in a defined medium for syntrophic growth and validating whether Mm can produce methane in a Dv-dependent manner. *In silico* flux predictions were performed using the COBRA -The COnstraint-Based Reconstruction and Analysis toolbox “optimizeCbModel” function and “fluxVariability” function in MATLAB. All model simulations related to FBA were performed on the MATLAB_R2019a platform using the recent version of COBRA. State-specific models were analyzed using custom scripts to remove highly variable reactions (Supplementary Figure 2 and Supplementary Table 2) and reaction fluxes were normalized per day by calculating the log2 fold-change differences^37^ in planktonic to sediment fluxes.

### Data availability

The DNA and RNA datasets generated during the current study are available under NCBI BioProject PRJNA974067.

## RESULTS

### Adaptation of a laboratory evolved syntrophic community to a simulated sub-surface environment

Custom anaerobic FBRs were designed as an analog for a subsurface environment, wherein sediment and groundwater are in flux and free to exchange metabolites and biota. Three independent FBRs were inoculated concomitantly with a minimal assemblage of the Dv-Mm SynCom obtained through end-point dilution of the mutualistic community adapted to obligate syntrophy conditions through 1,000 generations of laboratory evolution^2–4^. Each FBR contained 75 g of sediment (210 - 297 μm crushed quartz) and approximately 350 mL (FBR and reservoir) of lactate medium without sulfate to impose obligate requirement of growth by syntrophy (**Table 1** and **Fig. 1A**). The sediment bed in the FBRs were briefly fluidized by recirculating the lactate medium from the bottom of the reactor and up vertically through the column to mix cells throughout the reactor. The SynCom was cultured in batch mode for approximately 48 hours, as done previously^18^. After 2 days of growth, the FBRs were set to continuous fluidization (350 mL ⋅ min^-1^) with a duty cycle of 1 hour on and 1 hour off for the remainder of the experiment. The triplicate reactors were operated anaerobically (DO ∼1 mg/L) at pH ∼7.2 for approximately 175 hours under semi-continuous batch culture conditions with supplementation of fresh lactate medium at 7.8 mL ⋅ h^-1^, with a net dilution rate of 0.022 h^-1^ (**Table 1, Sup. Fig. 1**), which was similar to conditions used for growing biofilms (0.017 h^-1^)^18^.

**Table 1.** Operational parameters for anaerobic FBR arrays (n = 3).

| Operational Parameters | Settings | Notes |
| --- | --- | --- |
| Total Reactor Liquid Volume | ~350 mL | ~150 (body), 200 (reservoir), |
| Cross-sectional Area | 12.56 cm<sup>2</sup> |  |
| Pump Recycle Rate | ~350 mL · min<sup>-1</sup> |  |
| Volume Flux | 20.54 mL · min<sup>-1</sup> · cm<sup>-2</sup> |  |
| Influent Volumetric Flow Rate | 0.13 mL · min<sup>-1</sup> (7.8 mL · h<sup>-1</sup>) |  |
| Reactor Dilution Rate | 0.022 h<sup>-1</sup> | Flow Rate/Volume |
| Duty Cycle | 1:2 | 1 hour on: 2 hour period |
| Diffuser Plate (Frit Size) | 160 - 250 µm | Borosilicate Glass |
| Sediment Size | 210 - 297 µm | SiO<sub>2</sub> (Crushed Quartz) |
| Sediment Mass in Reactor | 75 g |  |
| Sediment density | 1.622 g · cm<sup>3</sup> |  |
| Reactor Tubing | Viton, Neoprene |  |

**Figure 1.**
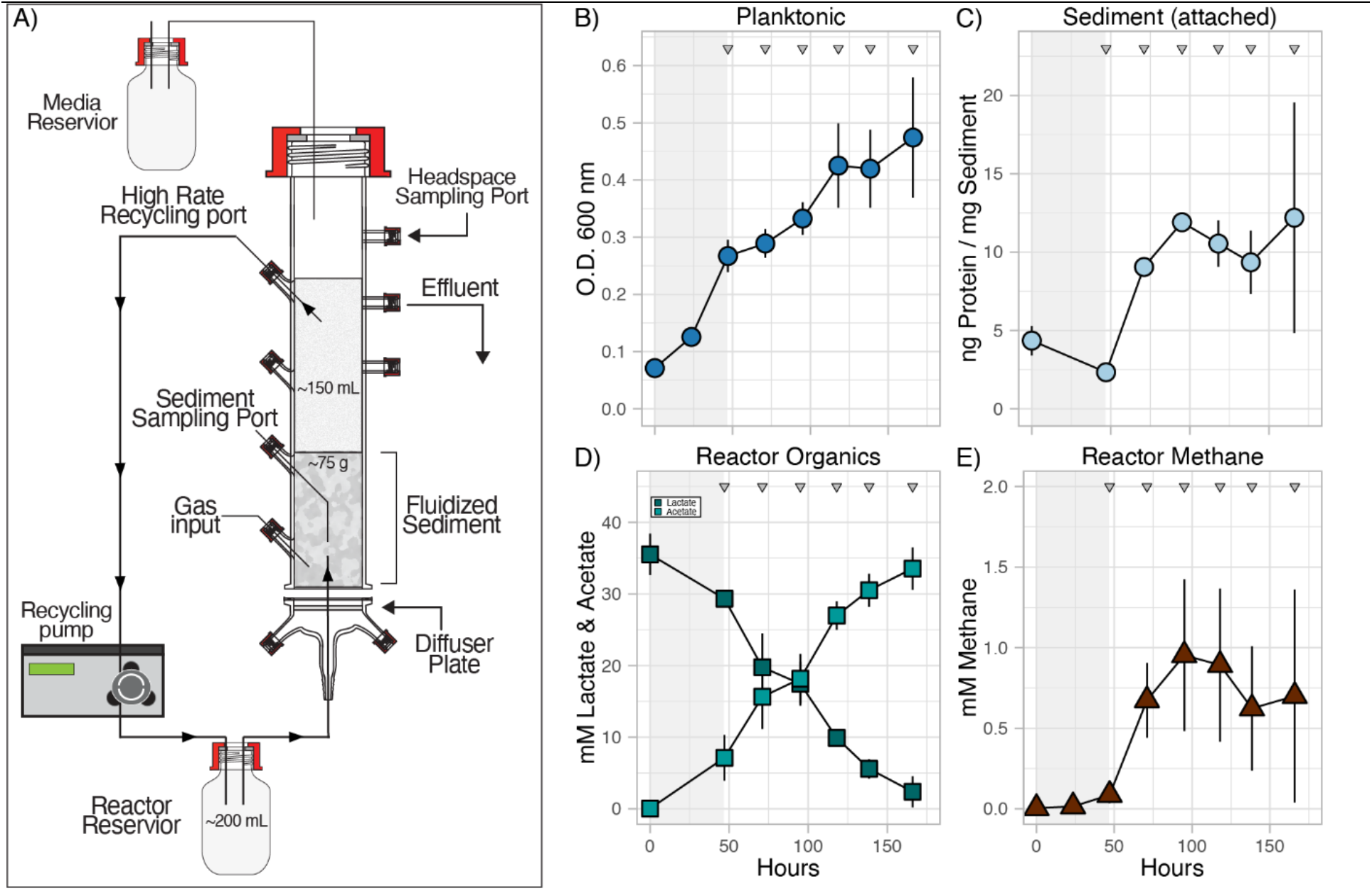
Partitioning of attached and planktonic syntrophic communities in FBRs. **A)** A schematic of the custom FBRs developed to simulate biphasic growth of microbial communities by recirculating growth medium upward through a column of sediment. Temporal growth dynamics of planktonic **(B)** and sediment attached **(C)** phases of the microbial community. Metabolite profiles provide confirmation syntrophic growth via the oxidation of lactate to acetate by Dv **(D)** and the production of methane by Mm **(E)**. The grey opaque background bars indicate the first 48 hours of batch growth in the FBRs prior to fluidization and the grey triangles represent the timepoints when DNA and RNA were sampled starting at day 1 to day 6. All error bars indicate standard deviation across 3 replicate FBRs.

Biomass in the planktonic phase increased at a steady rate with a maximum growth rate of 0.23 d^-1^ and an average doubling time of ∼3 days (**Fig. 1B)**. Using total protein from washed sediment as a proxy for biomass we determined that the attached community accumulated at a slower rate of 0.087 d^-1^ (doubling time: ∼8 days) (**Fig. 1C)**. Syntrophic coupling between Dv and Mm was evident in sustained lactate oxidation to acetate, with the subsequent production of methane (**Fig. 1D & 1E**). While significant increase in methane production was observed within 72 hours, the levels were steady at ∼0.75 mM for the remainder of the experiment. However, it should be noted that since the reactor headspace was allowed to vent through a one-way check valve to prevent pressure from exceeding 1 atm, the methane measured does not reflect the total amount produced throughout the experiment. Together, these results demonstrated that the co-culture established syntrophic growth in the simulated subsurface environment within the FBRs.

#### Heterogeneity of population structure and genetic variability across growth phases

Total DNA and RNA was sampled daily for 6 days once the reactor fluidization was turned on at approximately 48 hours (**Fig. 1. Grey triangles**) to evaluate population dynamics and functional heterogeneity of Dv and Mm across attached and planktonic growth phases. Based on normalized numbers of metagenomic sequencing reads corresponding to each organism, we determined that Dv was the dominant partner in both phases but community composition was significantly different between the planktonic and attached communities (**Fig. 2A**). For instance, Mm in the planktonic phase nearly doubled in relative abundance from day 1 to day 2 but thereafter accounted for about a third of the SynCom composition (i.e., ∼3:1 Dv:Mm). In striking contrast, the relative proportion of Dv in the attached SynCom was ∼20 times that of Mm. The relatively lower abundance of Mm in the attached SynCom is likely because Dv attachment occurs first and acts as a scaffold for Mm to join the biofilm, which takes ∼25 days for full maturation^18^. However, it is also likely that variants of both organisms within the evolved SynCom have different propensities of adaptation to syntrophic interactions in either the planktonic or the attached phases. Regardless, these findings suggested different mechanisms of syntrophic interactions had likely manifested across the attached and planktonic phase SynComs.

**Figure 2:**
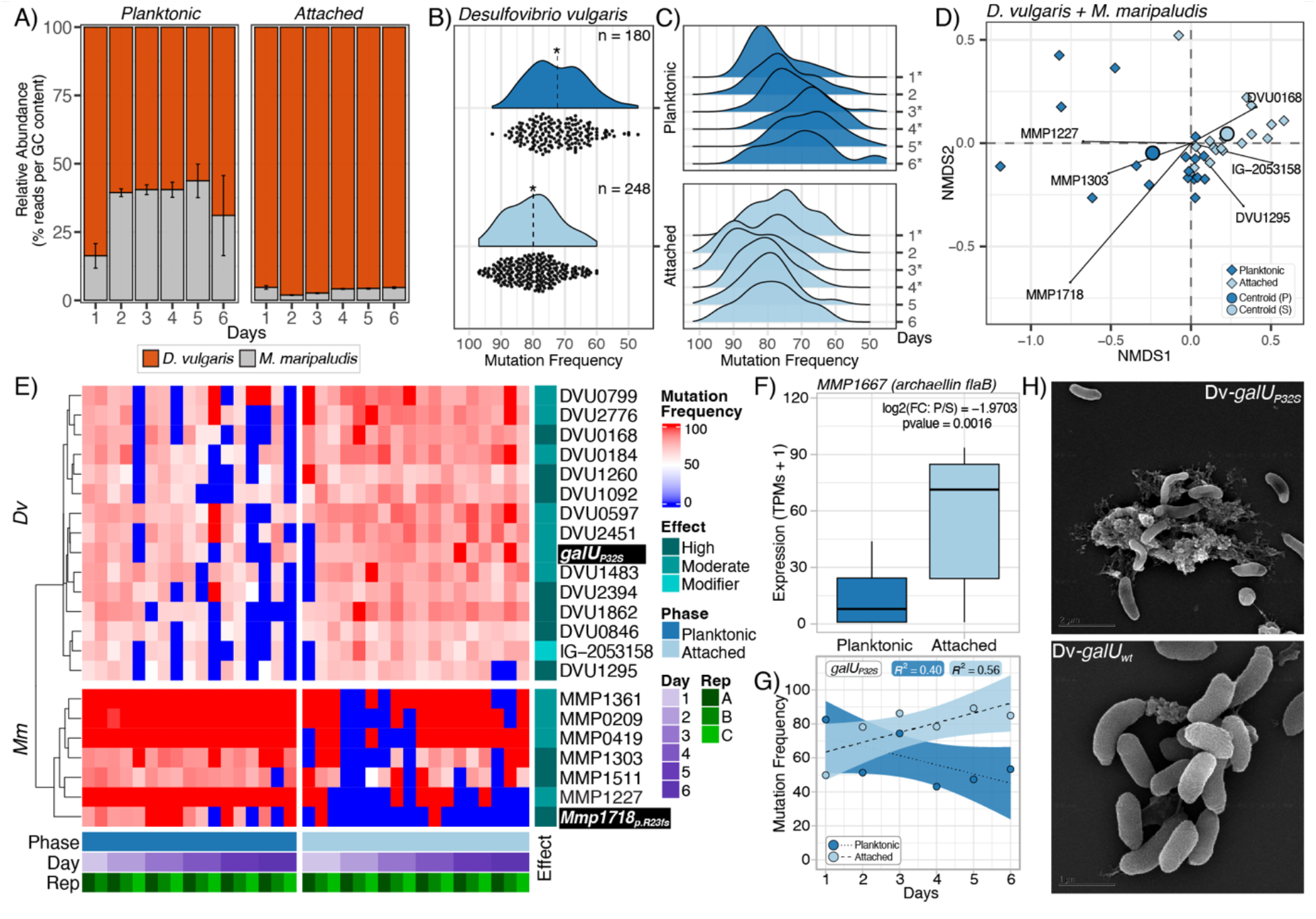
Population structure and dynamics between attached and planktonic communities. **A)** Longitudinal change in relative abundance of Dv and Mm, across the attached and planktonic communities, calculated from percent reads per GC content across all three reactors. **B)** Significantly different (p-value < 2.2e-16) distribution of Dv non-zero mutation frequencies between planktonic (mean 72.3%, dashed line) and attached populations (mean 79.9%, dashed line). **C)** Distribution of Dv non-zero mutation frequencies by day. Mutation frequency in planktonic populations significantly decreases over time, specifically on days 3, 4, 5, 6 (* designates a p-value < 0.05), while attached communities show stable or slight shift to higher frequency mutations. **D)** NMDS plotting of each sample (n = 35) for both Dv and Mm across phases. Arrows show ordination of specific mutations in NMDS space representing the optima habitat of mutation variant (showing top 3 for Dv and Mm). Large circles represent the centroid of the planktonic or attached community groupings. **E)** Heatmap of all 22 mutation variants clustered (Pearson correlation) across phases, days and replicate reactors. Selected variants are highlighted with a black background. **F)** Significantly decreased expression of MMP1667 (archaellin *flaB*) over days 3, 4, 5, 6 in the planktonic phase as a consequence of regulatory loss-of-function (*Mmp1718*_*p*.*R23fs*_) compared to the attached phase (*Mmp1718*_*wt*_ i.e., no loss-of-function). For boxplots center line, median; box limits, upper and lower quartiles; whiskers, 1.5x interquartile range; points, outliers. **G)** Mutation frequency of nonsynonymous variant in *galU*_*P32S*_ (DVU1283) highlighting the regression (R^2^ = 0.56) positive trend over time for the attached phase compared to planktonic. The ribbon on each linear regression plot represents the standard error. **H)** Scanning electron microscopy images showing overproduction of LPS by evolved Dv clones with the nonsynonymous *galU*_*P32S*_ mutation, but not by ancestral Dv-*galU*_*wt*_ cells.

We analyzed the metagenomic data with sequence alignment and variant calling (see Supplementary Methods) and analyzed frequency of Dv and Mm variants in SynComs across the two phases. The overall distribution of mutation frequencies of Dv was significantly lower in the planktonic as compared to the attached SynCom (**Fig. 2B**). Further analysis revealed that the frequency distributions of Dv mutations were statistically similar across the two phases on day 1, but over time it became progressively lower in the planktonic phase, and remained relatively stable in the attached phase (**Fig. 2C**). This progressive decrease in Dv mutation frequencies over time in the planktonic phase was likely due to wash out of genotypes with growth rates lower than the dilution rate (**Sup. Fig. 3A and B**). Surprisingly, the overall mutation frequencies of the Mm population remained constant in both phases. We performed nonmetric multidimensional scaling (NMDS) analysis to investigate if there was evidence of ecological differentiation^38^ by specific Dv and Mm variants (**Fig. 2D**). Strikingly, while most Dv variants, including DVU0168 and DVU1295, oriented towards attached habitats, most Mm variants, including MMP1227 and MMP1718, showed clear ordination towards planktonic growth (**Fig. 2D** shows change in abundance of top 3 variants of each Dv and Mm). Strikingly, the MMP1718 variant and was only observed 2 of 18 samples in the sediment phase (**Fig. 2E)**. Consistent with the demonstrated role of MMP1718 as a transcriptional activator of the *fla* operon ^39^, a loss-of-function frame-shift mutation at arginine 23 in this gene (*Mmp1718*_*p*.*R23fs*_) resulted in decreased expression of archaellum genes (specifically, *flaB2*) in the planktonic phase (**Fig. 2F**), which may have disrupted surface attachment of Mm^40,41^. While this was also consistent with the observation that the Mm-*Mmp1718*_*p*.*R23fs*_ variant had likely enhanced fitness during laboratory evolution, which was performed through sequential transfers of the planktonic phase evolved co-cultures^4^, it was intriguing that the WT allele “stuck around” even after end point dilution and ultimately swept through the Mm population in the attached phase.

**Figure 3.**
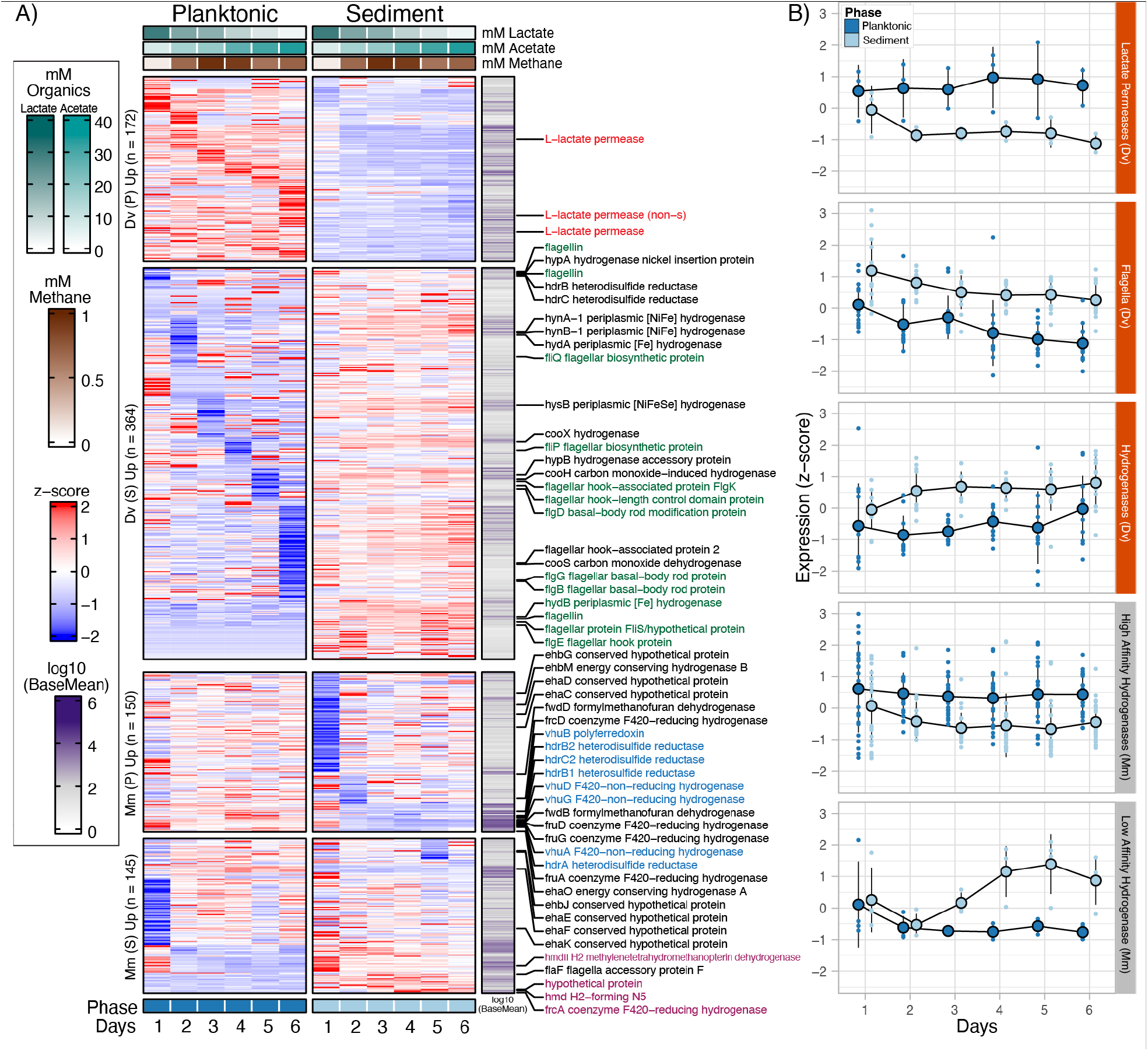
Transcriptome profiling shows distinct expression responses of co-existing planktonic and attached syntrophic communities. **A)** Heatmap (rows: genes) of z-scored expression (TPMs) of sequentially ordered significantly expressed genes (Fold Change > 2, p-value < 0.01) for each day (planktonic vs attached) and split between up- or downregulated genes for Mm and Dv. Right side annotation represents the log_10_ mean expression level for each gene across all samples (n = 35). **B)** Genes encoding central functions for syntrophic interactions between Dv (orange) and Mm (gray) have distinct expression patterns across planktonic and the sediment attached phase.

Of the total 22 (15 Dv and 7 Mm) mutations observed, nearly half (n = 10) were high impact mutations, meaning they had a gain or loss of start and stop codons or frameshift mutations. The other mutations that introduced nonsynonymous change, deletion or insertion of codon(s) were of moderate impact (**Fig. 2E**). Using linear regression across the three independent reactors (**Sup. Fig. 3B**), we discovered that while the frequencies of most Dv variants were stable in the attached community, mutations in a few Dv genes showed preference for the attached phase. In particular, a nonsynonymous variant that changed a non-polar proline to a polar serine in a UTP-glucose-1-phosphate uridylyltransferase (i.e., *galU*_*P32S*_), increased in frequency in the attached phase (R^2^ = 0.56) (**Fig. 2G**). Further investigation, using scanning electron microscopy, demonstrated that the nonsynonymous mutations in *galU* had likely resulted in lipopolysaccharide overproduction that may have facilitated attachment to sediment (**Fig. 2H**)^42–44^. Thus, the NMDS analysis uncovered evidence that specific variants of both organisms had likely led to niche differentiation of the SynCom ecotypes across the planktonic and attached phases.

#### Distinct mechanisms of adaptation and mutualism manifest across co-existing planktonic and attached syntrophic communities

We analyzed longitudinal transcriptome changes to understand adaptive consequences of ecological differentiation on the nature of syntrophic interactions across the planktonic and attached communities. A similar proportion of genes in Dv (536 genes, 14% of total genes), as compared to Mm (295 genes, 17% of total genes), were differentially expressed over time and across the two phases. There was a cascading pattern of changes in gene expression across the planktonic phase cells of Dv, but the transcription of these genes was relatively stable with sustained up- or downregulation in the attached phase cells (**Fig. 3A**). These expression patterns of Dv genes likely reflect adaptation of a stable, slow growing, sessile community to the attached phase, as compared to faster growing planktonic cells that experience greater environmental dynamics. In stark contrast, although Mm transcriptional response was relatively uniform over time in both phases, there was significant shift in expression levels of 173 genes during transition from day 1 to day 2 (**Fig. 3A**), which is consistent with how the two organisms establish syntrophy, especially when attaching to surfaces, with SRB colonizing first and subsequently laying the foundation for the methanogen to attach^18^.

Expression patterns of key genes for motility and attachment, lactate utilization, and methanogenesis across the organisms revealed a striking picture of distinct physiological states underlying interspecies interactions across the two phases. For instance, expression of 14 flagellar biosynthesis genes in Dv were constitutively down- or upregulated in the planktonic and attached phases, respectively, which was consistent with the known role of these genes in driving attachment^40,41^ and partner selection^45,46^. A similar expression pattern was observed for Mm, although in this case, the constitutive downregulation of the archaellum encoding genes could be attributed to selection of loss-of-function mutations in the cognate transcriptional activator (i.e., *Mmp1718*_*p*.*R23fs*_) in the planktonic phase. Further, all three lactate permeases in Dv were upregulated in the planktonic phase, especially day 4 onwards when there was an accumulation of biomass and a decline of available lactate for oxidation, which is the primary mechanism for generating H_2_ as a reductant for establishing syntrophic growth. Interestingly, upregulation of DVU2451, which can functionally compensate^47^ for the primary lactate permeases DVU2110 and DVU3026^48^, was likely a lactate scavenging mechanism by the planktonic Dv cells. By contrast, the attached Dv cells appeared to be not lactate limited as all permeases were constitutively downregulated in sediment.

We also observed distinct expression patterns of methanogenesis genes across the two phases, with some genes upregulated in the planktonic phase (e.g., formylmethanofuran dehydrogenase and F420-reducing hydrogenase) and others upregulated in the attached phase (e.g., H_2_ methylenetetrahydromethanopterin dehydrogenase). Notably, while the low affinity hydrogenases^49^ in Mm were upregulated in the attached phase, the high affinity hydrogenases were upregulated in the planktonic phase. This finding is consistent with the expectation of significantly higher H_2_ partial pressure in the microenvironment surrounding Mm in the attached phase, due to physical proximity to Dv cells, which were also in relatively higher abundance (Dv:Mm ∼18:1) in sediment. Conversely, lower cell density and relative abundance of Dv likely generated lower H_2_ partial pressure in the planktonic phase, triggering upregulation of the high affinity isoform of the hydrogenase in Mm as a mechanism for H_2_ scavenging (**Fig. 3B**). Thus, results of the expression analysis illustrated how ecological differentiation driven by differential segregation of mutants (e.g., Dv-*galU*_*P32S*_ and Mm-*Mmp1718*_*p*.*R23fs*_) led to further niche partitioning through differential expression of key genes across the two phases of growth within the same reactor, while still maintaining nutritional cooperation (i.e., syntrophy).

### Flux balance analysis using a curated genome-scale metabolic model for syntrophy reveals differential reaction fluxes across the two phases

A genome-scale metabolic network model of syntrophy was extended and curated to investigate reaction flux through the two members of the syntrophic community. In brief, metabolic models iJF477 (Dv)^33^ and iMR539 (Mm)^34^ were updated by curating 1014 and 688 reactions (see Supplementary Methods), respectively, and subsequently integrated into a genome-scale model of syntrophic interactions by setting interfacial constraints for interspecies exchange of 54 metabolites, including formate, H_2_, and alanine^19,21^. Further, the new syntrophy model (iSI1283) was contextualized to planktonic and attached phases of growth by using the GIMME algorithm^35^ to curate reactions based on longitudinal transcriptome profiles of the two organisms across six days of adaptation to two modes of growth across three reactors (**Fig. 4A, Supplementary Methods, Table 2**).

**Table 2.** Summary of metabolic models.

| Model | System | Reactions* | Genes | Metabolites |
| --- | --- | --- | --- | --- |
| iJF744 | Dv strain Hildenborough | 1014 (1016) | 744 | 948 (951) |
| iMR539 | Mm strain S2 | 688 | 539 | 710 |
| iSI1283 | Dv-Mm SynCom | 1707 | 1283 | 1658 |
| GIMME-contextualized<br>models | Day 1 planktonic | 1528 [924] | 1076 [592] | 1491 [892] |
|  | Day 1 sediment | 1578 [956] | 1108 [617] | 1553 [921] |
|  | Day 2 planktonic | 1616 [959] | 1111 [606] | 1592 [925] |
|  | Day 2 sediment | 1611 [946] | 1121 [610] | 1594 [912] |
|  | Day 3 planktonic | 1585 [931] | 1105 [604] | 1582 [920] |
|  | Day 3 sediment | 1598 [952] | 1121 [609] | 1573 [922] |
|  | Day 4 planktonic | 1621 [957] | 1115 [601] | 1598 [924] |
|  | Day 4 sediment | 1632 [966] | 1121 [615] | 1611 [928] |
|  | Day 5 planktonic | 1603 [954] | 1098 [601] | 1585 [926] |
|  | Day 5 sediment | 1617 [950] | 1124 [611] | 1599 [916] |
|  | Day 6 planktonic | 1587 [924] | 1103 [593] | 1577 [901] |
|  | Day 6 sediment | 1587 [924] | 1103 [593] | 1577 [901] |

\*The numbers within the ( ) parentheses indicate uncurated model and the numbers within the “[ ]” indicate counts specific to Dv organism

**Figure 4.**
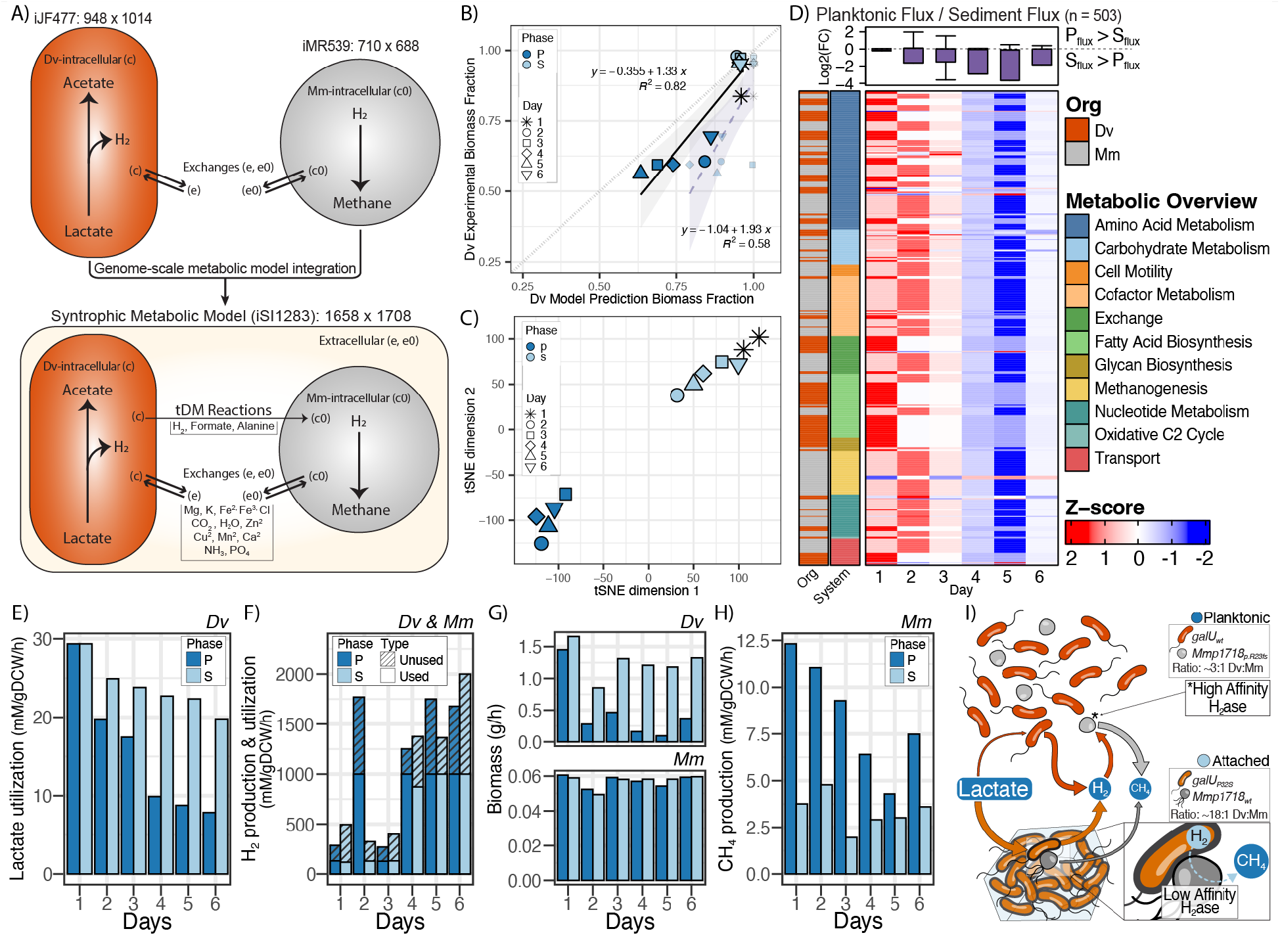
An integrated multi-species metabolic model reveals distinct metabolic flux states across attached and planktonic phases of syntrophic growth. **A)** Schematic representation of a new syntrophic metabolic model (iSI1283) obtained through the integration of genome-scale metabolic networks of iJF477 and iMR539. The syntrophic model highlights the general exchange of key metabolites for hydrogenotrophic methanogenesis and the required *in vivo* exchanges, transports and reactions from intracellular to extracellular. **B)** *In silico* prediction of Dv to Mm proportions before (small, transparent points, with dashed purple line) and after (large, solid points, with solid black line) bi-level optimization. The ribbon on each linear regression plot represents the standard error. **C)** t-SNE analysis of longitudinal differential flux states of Dv and Mm across planktonic and attached phases. **D)** Boxplots show distribution of log2 fold-change of flux in planktonic phase as compared to sediment phase for differential active reactions (n = 503) (**See Supplementary Table 2**) with biomass as the objective function. Heatmap shows row z-scores for each relative flux change between the two phases and across all days. Using the FBA model with methane production as the objective function, we predicted the individual fluxes of key syntrophic methanogenesis metabolites for each phase and day; **(E)** Dv lactate, **(F)** Dv H_2_ production (Total bars: hashed and non-hashed), Mm H_2_ utilization (non-hashed), **(G)** the predicted flux towards biomass for each organism, and **(H)** the amount of methane produced by Mm. **H)** A heuristic model for the predicted differences in flux between relative ratios planktonic and attached cells and ecotypes (Dv-*galU*_*wt*_ —Dark orange: *Mmp1718*_*p*.*R23fs*_ —Light grey and no archaellum; Dv-*galU*_*P32S*_ —Light orange and thick membrane: *Mmp1718*_*wt*_ —grey with archaellum) The weight of each arrow reflects the amount of flux predicted through or to each organism and phase.

Notably, accurate prediction of significantly higher relative abundance of Dv to Mm in the attached as compared to the planktonic phase (**Fig. 4B**) demonstrated that the contextualized iSI1283 models had captured distinct mechanisms by which the SynCom had partitioned between the two phases. For further investigation of these evolved strategies, the experimentally quantified relative abundances of Dv and Mm (**Fig. 2A**) were included as constraints for bi-level optimization of syntrophic growth of both organisms, which significantly improved model accuracy (R^2^ = 0.82 from 0.58) (**Fig. 4B**). Further, t-SNE (Stochastic Neighbor Embedding) analysis with bi-level optimized models demonstrated that while flux states of the SynCom were indistinguishable on day 1, they became increasingly divergent over time as the communities partitioned across the two phases (**Fig. 4C**), with significantly greater overall reaction flux across multiple pathways in the attached phase (**Fig. 4D; boxplots**). Strikingly, the models predicted a dramatic metabolic shift upon transition from day 3 to 4, driven by increased flux through metabolic pathways within the attached cells (**Fig. 4D; heatmap**). The timing of the metabolic shift coincided with an inflection point when concentration of acetate in the reactors exceeded corresponding instantaneous levels of lactate in the reactor (**Fig. 1D**).

Using methane as the objective function, we used the iSI1283 models to uncover relative contributions of SynComs in the attached and planktonic phases to overall productivity of hydrogenotrophic methanogenesis in the FBRs. This analysis predicted that while rate of lactate utilization by Dv was similar across both phases on day 1, it declined at a significantly faster rate in the planktonic phase. By day 3 the SynCom in the attached phase utilized lactate at a rate that was twice that of planktonic cells (**Fig. 4E**). However, despite the high rate of lactate utilization by Dv in sediment, the rates of hydrogen (H_2_) production and consumption by the SynComs (**Fig. 4F**) were similar across the two growth phases. After day 3, the production rate of H_2_ (Dv) exceeded the maximum amount needed (1,000 mM gDCW^-1^ h^-1^; DCW: Dry Cell Weight) to support predicted increases in biomass of both planktonic and attached SynComs (**Fig. 4G)**. Methane production was significantly higher in the planktonic phase, likely due to the excess H_2_ from both phases and the higher relative proportion of Mm (**Fig. 4H)**. Interestingly, the model predicted that a smaller fraction of the H_2_ captured by the low affinity hydrogenases was utilized for methane production by Mm in the attached community. The excess H_2_ from the attached community was released into the growth media and likely scavenged by high affinity hydrogenases towards significantly higher rate of methanogenesis by planktonic Mm (**Fig. 4I**).

## DISCUSSION

Our study demonstrates that, even within a simple two organism community that originated from clonal isolates, ecotypes adapted to different biogeographies can arise very rapidly, introducing complex structure and division of labor, thereby increasing the overall productivity of the community. In the soil subsurface, microbial communities can be more productive in attached (sediment) relative to planktonic phase^40,41,50^. Two explanations have been put forth to explain this phenomenon: first, that the particulate material in sediment is largely organic and nutrient rich, or inorganic material with adsorbed nutrients^41,51,52^. Second, it has been argued that an attached community in a flowing environment has greater access to soluble nutrients due to change in the depth of the diffusive boundary layer - the greater the flow, the thinner the layer and greater the flux of nutrients through the boundary layer^40,53–55^. Both of these mechanisms may be at play in the FBRs and responsible for higher rate of lactate utilization by Dv in sediment, also explaining why the Dv-*galU*_*P32S*_ mutant might have gained fitness advantage through attachment. By contrast, Dv cells in the planktonic phase likely went with the flow and were therefore relatively less efficient at accessing and utilizing lactate, although they leveraged their diverse physiological capabilities as generalists by continually adapting to changing availability of nutritional sources as reflected in the cascade of changes in expression of large numbers of genes. The high amounts of H_2_ produced by Dv generated a higher local H_2_ partial pressure in sediment, however H_2_ was not completely utilized by attached Mm cells despite their physical proximity. Attached Mm cells likely upregulated low affinity hydrogenases to aid H_2_ consumption and reduce H_2_ partial pressure in the microenvironment of attached cells to prevent feedback growth inhibition of Dv. Interestingly, in the attached phase, Mm was also less efficient at transforming H_2_ to methane, and as a result excess H_2_ was released into the fluid environment, where planktonic Mm used high affinity hydrogenases to capture/scavenge H_2_ and use it towards copious amounts of methane production. Indeed, Mm existed in higher relative proportion to Dv in the planktonic phase as compared to sediment, and this also explains why the loss-of-function mutation in the archaellum regulator (*Mmp1718* _*p*.*R23fs*_) conferred higher fitness in the planktonic phase (**Fig. 4I**). In other words, despite the mixing of genotypes, ecotypes that had differentially adapted to sediment and planktonic phases segregated into distinct communities that were functionally interdependent.

It is clear that interplay of ecotypes of the two organisms across the planktonic and attached phases greatly improved cooperativity and overall productivity (i.e., biomass and methane production) of the entire mutualistic community^4^. But the emergence of a sediment adapted evolved ecotype in this interplay was perplexing because the laboratory evolution was specifically designed for selection of mutations that optimized syntrophy in a planktonic growth state through serial transfers of liquid co-culture. In other words, it is intriguing how mutations that improved characteristics of the attached community arose and how they “stuck around”. For instance, it is understandable that the *Mmp1718*_*p*.*R23fs*_ mutation was selected because it promoted better planktonic growth of the methanogen, but it’s puzzling how the ancestral allele persisted in the population at low frequency, even after end point dilution. Our results show that the ancestral *Mmp1718*_*wt*_ allele may have benefited from improved physical interactions with Dv, especially upon appearance of the lipopolysaccharide overproducing *galU*_*P32S*_ mutant. The Dv-*galU*_*P32S*_ mutant may have segregated with Mm-*Mmp1718*_*wt*_ as attached communities on the walls of balch tubes, which begs the question of how they persisted despite multiple serial transfers of the liquid culture, especially given that establishment of syntrophy in a biofilm takes significantly longer than the period of culturing between transfers^18^. Perhaps, the Dv-*galU*_*P32S*_ and Mm-*Mmp1718*_*wt*_ variants survived as free floating cell-to-cell interactions (i.e., attached partners)^45^. Regardless, our findings suggest that the planktonic and attached lifestyles of the two organisms in a syntrophic association are intricately intertwined such that optimization of one lifestyle drives co-optimization of the other. This hypothesis is supported by our findings that sediment attachment of the Dv*-galU*_*P32S*_ mutant was associated with improved lactate utilization, and may have required physical interactions with Mm-*Mmp1718*_*wt*_ for reducing H_2_ partial pressure in its microenvironment, and conversely, improved planktonic growth of the Mm-*Mmp1718*_*p*.*R23fs*_ mutant was associated with more efficient H_2_ utilization and also likely benefitted from metabolite exchange with free Dv cells that continually adapted to changing conditions of the flowing medium. This interplay of physiologies associated with planktonic and attached ecotypes of the two organisms is likely ingrained in the molecular networks of the two organisms, such that mutations affecting one physiological state potentiated selection of mutations in genes associated with the other linked physiological state.

Our results demonstrate that physical segregation of specific genotypes that arose over short evolutionary timescales led to ecotype differentiation and resource partitioning wherein the SynComs manifested distinct physiological states and vastly different community structures across the attached and planktonic phases. Yet there was meaningful interplay between the attached and planktonic communities that likely contributed to overall higher productivity. Thus our study findings shed insight into the evolutionary mechanisms by which microbial communities leverage ecotype differentiation to divide and conquer spatiotemporally complex environments. Remarkably, our findings also show that ecotype differentiation occurred within 300-1,000 generations by mutations in as few as 15 genes in Dv and 7 in Mm, which is consistent with observations in natural microbial communities in the environment^15,56,57^. Importantly, all ecotype differentiating mutations in the SRB and the methanogen, respectively, had high G-scores, which is a measure of reproducibility of selecting mutations in the same gene across many independent evolution lines^4^. High G-scores indicates that the ecotype differentiating mutations (including the *Mmp1718*_*p*.*R23fs*_ and the *galU*_*P32S*_) were observed independently and at high frequency across multiple evolution lines demonstrating further that this mechanism of ecotype differentiation was fairly common and also that the attached community (primarily Mm-*Mmp1718*_*wt*_ and Dv-*galU*_*P32S*_) had co-optimized with the planktonic community across multiple lines. We predict that prolonged laboratory evolution of such a microbial community with co-existing but physically segregated genotypes that exclude each other across habitats could likely lead to sympatric speciation, wherein mutually excluding ecotypes of the same species diverge to a point where horizontal exchange of genetic material through homologous recombination is no longer feasible^6,14,15,58^.

## ACKNOWLEDGEMENTS

This material by ENIGMA- Ecosystems and Networks Integrated with Genes and Molecular Assemblies (http://enigma.lbl.gov), a Science Focus Area Program at Lawrence Berkeley National Laboratory is based upon work supported by the U.S. Department of Energy, Office of Science, Office of Biological & Environmental Research under contract number DE-AC02-05CH11231.

## AUTHOR CONTRIBUTIONS

J.J.V. designed custom anaerobic fluidized bed reactors and designed experiments with S.T., and N.S.B. J.J.V., J.W., and M.R. performed all growth experiments, extractions (DNA, RNA, and protein). M.R. and K.A.H. performed all IC an GC analysis. M.A. and M.Z. performed SEM analysis on variant strains. S.T. performed variant calling, RNA alignment, and transcript quantification, while J.J.V. analyzed the mutational output files and performed differential expression analysis. J.J.V. and S.R.C.I. integrated and refined the iSI1283 model and conducted additional analyses. J.J.V., S.R.C.I., S.T., S.M.G., K.A.H., D.A.S., and N.S.B. all contributed to the discussion of results and conclusions. J.J.V., S.R.C.I., and N.S.B wrote the manuscript.

