## Supplementary Information for "Origin of biogeographically distinct ecotypes during laboratory evolution"

**Supplementary Methods.** Supplementary Materials/Subjects and Methods.

**Supplementary Figure 1. Longitudinal profiles of dissolved oxygen and pH from reactor medium.** Dissolved oxygen and pH (DO [1 mg/L = 1 ppm], green diamonds) from the recirculating media inside triplicate FBRs was measured daily using benchtop probes. The grey opaque bars indicate the first 48 hours of batch growth in the FBRs prior to fluidization and the grey triangles represent the timepoints when DNA and RNA were sampled. All error bars indicate standard deviation across 3 replicate FBRs (n = 3).

**Supplementary Figure 2. Distribution of FBA reaction fluxes across days and phases and selected cutoffs to reduce complexity and increase confidence. A)** The variance in flux values (mM/gDCW/h) was compared to the standard deviation (Std) for each reaction across each day and phase (n = 12) to identify a confidence threshold (2 Std or 95% confidence intervals, dashed line) for removing highly variable reaction fluxes**. B)** Reactions fluxes below a Std. of 2 that remain after filtering. **C)** Histogram of the non-variable reaction fluxes across all days and phases indicating most reactions fluxes are centered around zero **(D)**.

**Supplementary Figure 3. Regression analysis of Dv mutation frequencies over time for each phase. A)** Comparisons of each linear model’s coefficient of determination ($R^2$) with its slope for each variant at each phase. Labels were placed on each model that had an $R^2$ greater than .5 and DVU1283 ($galU_{P32S}$). **B)** Each regression model with $R^2$ values for all Dv variants across both phases with ribbons representing the standard error.

**Supplementary Table 1.** Summary of monitored parameters over the experimental time-course for the triplicate FBRs.

**Supplementary Table 2.** Workflow for analysis of differential flux states between planktonic and attached communities.

**Supplementary Data File 1.** Workbook containing the output analysis for the identification of mutations of the evolved cocultures and a short summary of the mutational impact and gene descriptions of the 22 variants.

**Supplementary Data File 2.** Workbook containing all differential expression analyses for each condition comparison for both Dv and Mm, including per day comparisons and across all days. The last sheet contains all significantly differentially expressed genes used to create Figure 3A.

**Supplementary Data File 3.** Workbook containing detailed spreadsheets of the iSI1283 constraint-based metabolic model and *in silico* flux predictions for the contextualized models of planktonic and sediment phases of the SynComs from Day 1 to Day 6.

### SUPPLEMENTARY METHODS

#### Generation of end-point dilutions[1] (Hillesland et al., 2014).

In this study, we used a simplified community of *Desulfovibrio vulgaris Hildenborough* (Dv) and *Methanococcus maripaludis* (Mm) derived from the end-point dilutions of cocultures that evolved together over 1,000 generations of syntrophic growth as previously described[1,2]. Briefly, two clones of Dv and Mm were combined to create 24 ancestral lines in coculture medium A (CCMA)[3] under anaerobic conditions (80% $N_2$:20% $CO_2$ headspace) in Balch tubes. Every week cocultures were transferred into a fresh media at a 100-fold dilution and incubated with or without shaking. This repeated propagation was continued for 152 weeks and populations were archived as frozen glycerol stocks after generations 100, 300, 500, 780, and 1,000 generations. Biomass collection was done as described before[1]. From the 1,000 generation glycerol stocks, end-point dilutions were generated by allowing cocultures to grow to stationary phase and then re-supplemented with fresh CCMA medium and incubated until cells reached late-exponential phase, which was then used to make two 10,000-fold diluted cultures in fresh CCMA media. The diluted cultures were then put through 10-fold serial dilutions and monitored for growth. This was followed by a second round of dilutions to finalize the simplified communities and made into glycerol stocks and stored at -80 °C.

#### Fluidized bed reactor setup and operation.

We designed and constructed custom glass anaerobic fluidized bed reactors operated by recirculating media from the bottom of the reactor to the top creating a vertical velocity through the column to create a bed of fluidized sediment. Three replicate FBRs were prepared with 75 g of 210 - 297 µm crushed quartz (Sigma-Aldrich: 50-70 mesh quartz) to act as the sediment bed sitting above a diffuser plate (P0 Frit: 160 - 250 µm), which allows the passage of liquid but not sediment. Reactors were autoclaved prior to inoculation and filled with sterile anaerobic lactate media (no sulfate) to an operating volume of approximately 350 mL. All three reactors were operated at the same time in a heat controlled room held at ~30°C. During operation of the FBRs the recirculating peristaltic pumps were set at 350 mL · $min^{-1}$ to a duty cycle of 1 hour on per 2 hour period. Media was supplemented with fresh lactate media using peristaltic pumps set to 0.13 mL · $min^{-1}$ and a duty cycle of 1 hour on per 2 hour period matched with the fluidization pumps. All tubing used during operation of the FBRs was Viton or Neoprene.

**FBR inoculation and sample analysis**.

A simplified SynCom from an end-point dilution of Dv and Mm (EPD9) was sourced from the previously described study[1,2]. Revived glycerol stocks were grown in anaerobic Balch tubes containing lactate media for three successive transfers to reestablish growth and then transferred to a 60 mL serum bottle for inoculation into FBRs. A 15 mL aliquot of optical density ($OD_{600}$) 0.3 culture from a single serum bottle was used to inoculate each FBR containing anaerobic lactate media (**Table 1**). FBRs were briefly fluidized to mix cells throughout the reactor and allowed to grow in batch mode for approximately 48 hours, as done in previous studies[4]. After approximately 48 hours of growth, the FBRs were switched to fluidization mode for the rest of the experiment.

While reactors were fluidizing, approximately 15mL of planktonic culture was sampled anaerobically using a sterile 20 mL syringe and needle. Planktonic fractions were split for $OD_{600}$ analysis (1 mL), protein extractions (2 mL), nucleic acid extractions (4 mL), nutrient analysis, pH (2 mL), and dissolved oxygen (DO) (2 mL). Protein and nucleic acid samples were kept on ice until pelleted. In a microcentrifuge tubes, 2 mL of planktonic culture was spun down and pelleted at 12,000 × g for 10 min at 4°C. The decanted supernatant was saved (-20°C) for organic acid (lactate and acetate) analyses, while the cell pellets for protein and nucleic acids were flash frozen in liquid $N_2$ and stored at -80°C until extraction. Lactate and acetate samples were diluted with millQ H2O and measured using a ion chromatography as described previously[5].

To sample the communities attached to sediment, we created sample chambers made from Balch tubes that were filled with 100% $N_2$. While reactors were fluidizing, sediment was sampled anaerobically by pulling a vacuum from the sampling port of the FBR into a sterile anaerobic Balch tube that was on ice. Immediately following sediment sampling from the FBR, samples (on ice) were taken directly to a 4°C cold room for washing. The sediment samples transferred to 15 mL conical tubes and planktonic culture was decanted into a waste container. Approximately, 10 mL of pre-chilled PBS (4°C) was added to the conical tubes and sediment was washed for 5 minutes on a benchtop tube rotator. The PBS wash solution was then decanted from the sediment samples and the 5 minute wash step was repeated with fresh chilled PBS. After the second wash, sediment samples were split into 2 mL microcentrifuge tubes (Protein, DNA, and RNA) and wash solution was pipetted out and samples were immediately frozen in liquid $N_2$.

We prepared glass vials filled with 100% $N_2$ and crimped closed with rubber septum and aluminum seal to store sampled headspace from the FBRs to be analyzed via gas chromatography (GC). Immediately prior to sampling reactor headspace, 2 mL of 100% $N_2$ was removed from the ethanol washed storage vials using a small diameter needle and 2 mL syringe. The same needle and syringe was then used to sample 2 mL of reactor headspace through the ethanol sterilized headspace sampling port and reinjected into the storage glass vials for quantification of methane using GC. Methane was measured using an SRI 8610C GC (SRI Instruments, Torrance, CA) equipped with a 6' silica gel column (SRI Instruments, Torrance, CA) at an oven temperature of 80°C and a flame ionization detector operated at 385°C. The carrier gas consisted of >99.998% $N_2$ gas (Praxair, Danbury, CT) at 20 mL · min$^{-1}$, >99.5% $H_2$ gas (Praxair, Danbury, CT) at 25 mL · min$^{-1}$, and air supplied via an internal pump at 250 mL · min$^{-1}$.

**DNA, RNA, and protein extractions.**

Nucleic acid samples for planktonic pellets and sediment samples were extracted using the MasterPure Complete DNA and RNA Purification Kit (Epicentre: #MC85200). Nucleic acid samples were then split between DNA and RNA samples. DNA samples were treated with RNase and RNA samples were treated with DNase. All samples were quantified using the Qubit DNA or RNA High Sensitivity (HS) kits (Invitrogen). Protein extractions for planktonic cell pellets were performed according to Thermo Scientific B-PER Complete Bacterial Protein Extraction Reagent protocol. For sediment samples, sediment was transferred to a 1.5 mL eppendorf tube and incubated with B-PER Bacterial Protein Extraction Reagent and rotated for 15 minutes to extract protein. After protein extraction, sediment was transferred to a glass petri dish and dried in a 70°C oven overnight to quantify mass of sediment. All protein samples were quantified using the Qubit Protein Broad Range Assay and normalized to dry sediment weight.

**DNA and RNA library preparations and sequencing.**

DNA samples were prepared for sequencing using the Nextera XT DNA library prep kit. Libraries were sequenced using a mid-output 2 x 75 bp for 150 bp fragments on a NextSeq 500. RNA samples were prepared for sequencing by performing rRNA depletion with a RiboZero Plus kit. Libraries were prepared using the Truseq Stranded Total RNA library prep kits, checked by Bioanalyzer (Agilent Technologies) high sensitivity RNA chips, and pooled for sequencing. Sequencing was performed using a mid-output 2 x 75 bp kit for 150 bp fragments using NextSeq 500. Custom scripts were used to quantify relative abundance from metagenomes of the SynCom based on percentage of reads per GC content for Dv (~63.2%) and Mm (~33.1 %) for each sample (n = 35).

**Identification of mutations.**

Mutations accumulated in populations were determined by using a custom sequence alignment and variant calling pipeline as described previously[6] (see also https://github.com/sturkarslan/evolution-of-syntrophy). This pipeline included quality control and trimming of the raw sequencing reads in fastq format by using Trim Galore software[7] (http://www.bioinformatics.babraham.ac.uk/projects/trim_galore). The alignment of the quality trimmed sequences to reference *D. vulgaris* (Genbank assembly: GCA_000195755.1.30) and *M. maripaludis* (Genbank assembly: GCA_000011585.1) genomes and subsequent processing steps before calling the variants was done by following The Genome Analysis Toolkit (GATK)[8] best practices. Briefly, reads were first aligned to the reference genome using Burrows-Wheeler Alignment Tool (bwa)[9] (version 0.7.17-r1188) in paired-end mode. The resulting alignment files in the SAM format were converted to BAM files, sorted and indexed by using Samtools version 1.9[10]. BAM files were marked for duplicates using Picard Tools (http://broadinstitute.github.io/picard/) (version 1.139), and local realignment around indels was performed to identify the most consistent placement of reads relative to the indels. Variant calling was performed independently by using three different algorithms including GATK UnifiedGenotyper, Varscan 2[11] (version 2.3.9) and bcftools from Samtools package. The default parameters were used for UnifiedGenotyper, whereas for Varscan parameters were --min-coverage 8 --min-reads2 2 --minavg-qual 30 and bcftools parameters were -vmO -s LOWQUAL -i'%QUAL>30. Variants identified by each caller were collated and filtered for variant frequency equal or greater than 20%. A variant was included in the analysis only if it is simultaneously called by at least two of the callers. The resulting variants were annotated using SnpEff tools[12] (version 4.3). Variant analysis, NMDS, and linear regressions were done using custom scripts and a list of compiled variants is described in **Supplementary Data File 1**.

**Scanning electron microscopy (SEM).**

Scanning electron microscopy (SEM) was used to visualize the overproduction of LPS using a 300 generation evolved line [1,2,6]. Monocultures were grown in 5 ml CCMA medium[3] supplemented with 10 mM sodium acetate and 30 psi $H_2$, whereas cocultures were grown in 20 ml CCMA medium supplemented with 30 mM sodium DL-lactate. Cocultures were incubated shaking at 37°C until reaching early (OD = 0.11-0.15) and late stationary phase (OD = 0.4-0.45). 1 ml of culture was removed, diluted to an OD of 0.08 and fixed with glutaraldehyde (1% V/V) and sent for SEM to the LBNL (Lawrence Berkeley National Laboratory).

Samples for SEM were prepared on 0.1% poly-l-lysine coated silicon wafers and a cell suspension was placed on the wafers for 15 minutes to allow cells to adhere. Excess cell suspension was then gently rinsed off with water and the wafers were fixed with 2.5% glutaraldehyde in Sodium Cacodylate (pH 7.2) for 1 hour and post-fixed with 1% Osmium Tetroxide in Sodium Cacodylate (pH 7.2) on ice for another hour and finally rinsed three times with Sodium Cacodylate buffer (pH 7.2) to remove the remaining fixatives. Dehydration was performed through a graded ethanol series (20%, 40%, 60%, 80%, 90%, 100%, 100%, 100% at 7 minutes per step), followed by critical point drying using the Tousimis AutoSamdri 815 Critical Point Dryer (Tousimis, MD, USA). After the samples were dried, they were sputter coated with gold-palladium using a Tousimis Sputter Coater (Tousimis, MD, USA) to prevent charging in the microscope. Images were collected using the Hitachi S5000 Scanning Electron Microscope (Hitachi High Technologies America Inc, CA, USA).

**Gene expression analysis.**

Paired-end Illumina reads were processed using Trim Galore[7] version 0.4.3 following Illumina default quality filtering steps. Reads were further trimmed for low-quality ends (Phred score <20) and cleaned up for adapter contamination with TrimGalore. For sequence alignment, reference genomic gene sets for *Desulfovibrio vulgaris Hildenborough* (ASM19575v1) and *Methanococcus maripaludis strain S2* (ASM1158v1) were combined to create a merged reference. Transcript abundance estimation was performed by using kallisto[13] v0.44.0. Differential gene expression analysis (**Supplementary Data File 2**) was performed using DESeq2 package[14] v1.22.2 in R after importing kallisto transcript abundance estimates as TPMs (Transcript Per Million) for each organism using the function "DESeqDataSetFromMatrix". Specifically, differential expression analysis was performed by comparing planktonic expression to the sediment expression for all comparisons using the "collapseReplicates" function from DESeq2 for each day separately (e.g.,

Day 1 Planktonic vs Day 1 Sediment) and for all days combined (Days 1 – 6 Planktonic vs Days 1 – 6 Sediment).

**Metabolic model refinement and integration.**

The genome-scale metabolic models of Dv (iJF744)[15] and Mm (iMR539)[16] were used in this study to represent Dv and Mm individually and further updated for accuracy. The published version of iJF744 metabolic model consisted of 1016 reactions and 951 metabolites. The gene-protein-reaction (GPR) relationship was represented using 744 genes of Dv in the network. These 1016 reactions were then curated as suggested in the publication. These updates included the renaming of the reaction 'rxn05938' as pyruvate synthase, removing the protein biosynthesis, DNA replication, and RNA transcription (i.e., reactions - 'rxn13782', 'rxn13783', and 'rxn13784') and fixing the capitalization of 'Ex_cpd00218' (Rxn name changed from 'Ex_cpd00218(e)' to 'EX_cpd00218(e)'). Two reactions ('rxn11934B_CC' 'QMO (Co-culture)' and 'rxn11934B_SR - 'QMO (Sulfate Reducing)') were added in the place of one reaction (rxn11934) and also the exchange reactions of hydrogen, acetate and pyruvate were renamed to 'EX_cpd11640(e)', 'EX_cpd00029(e)', 'EX_cpd00020(e)'. All of these updates were performed according to the 'alterDvhModel.m' script from the original publication. This curation of the iJF744 model resulted in 1014 reactions, 744 genes and 948 metabolites. We also updated the GPR of the reaction (rxn14403) as an "OR" relationship for the gene 208282 (DVU2776) because the sulfite reductase has been characterized to have multiple functions[17,18]. The published version of the iMR539 model consisted of 539 genes from Mm that were linked to a total of 668 reactions and 710 metabolites[16]. 51 exchange reactions in the Mm model were updated for the lower bounds to limit the metabolite exchanges for the syntrophic condition.

The integration of the two metabolic networks (iJF744 and iMR539) into a single syntrophy community model iSI1283 was achieved by interlinking the compartments of Dv and Mm through the known metabolite exchanges across Dv and Mm[19]. Initially, the metabolites of Mm were compartmentalized using standard tags for cytosol ("c0") and extracellular ("e0") compartments with '0' at the end to specifically mention that these metabolites belonged to Mm. Hence, these tags identify Mm as "Organism 0". Metabolite tags with ("c") and ("e") represent Dv. All the exchange reactions used for introducing metabolites to the extracellular compartment were standardized in "EX_{metabolite ID}[e0]" format. These exchanges were not removed because the ions and $co_2$ were allowed from extracellular i.e., the media. Thus, the iSI1283 model integrated the two networks (iJF744 and iMR539) directly, given that the metabolites and reaction ids are the same but contains these additional tags to identify them. All the metabolic model files are available for download from our GitHub repository https://github.com/baliga-lab/SynCom-model-for-DvH-and-Mmp and are produced in the **Supplementary Data File 3**.

**Flux predictions using state-specific models and analysis.**

The iSI1283 metabolic model was contextualized using gene expression data from planktonic and sediment phase obtained for day 1 to day 6. This step involved the use of the GIMME algorithm[20]. The GIMME algorithm takes the gene expression profile and reduces the model into a reduced network with all reactions that have genes expressed above the threshold. We applied the constraint-based method for simulating the metabolic steady-state of the SynCom model using flux-balance analysis (FBA)[20,21]. The initial validation steps involved checking the capacity of the SynCom model to produce biomass in a defined media for the syntrophic co-culture and checking whether Mm can produce methane in a Dv-dependent manner. *In silico* flux predictions were performed using the COBRA toolbox “optimizeCbModel” function and “fluxVariability” function in MATLAB. All model simulations related to FBA were performed on the MATLAB_R2019a platform using the recent version of COBRA -The COnstraint-Based Reconstruction and Analysis toolbox.

State-specific models were analyzed using custom scripts to remove highly variable reactions (Reaction fluxes < 2 standard deviations) and with flipped reaction signs prior to fold change analysis (**Supplementary Table 2**). 705 of the reactions had flux values of zero and were removed after reaction normalization by calculating the fold change of each reaction flux per day (e.g., Day 1 Planktonic vs Day 1 Sediment) as done for the gene expression analysis. This allowed for the comparison of fluxes between states resulting in a total reaction fold change matrix of 609 reactions. The reactions subsystems were further categorized by identifying the highest level of KEGG (Kyoto Encyclopedia of Genes and Genomes) associated metabolisms (https://www.genome.jp/kegg/pathway.html#global) to generate a “metabolic overview” tag.

### **Data and Code availability**.

***Data availability***. The DNA and RNA datasets generated during the current study are available under NCBI BioProject PRJNA974067. Links to the code used to analyze the data will be made public upon publication.

**Supplementary Figure 1**.

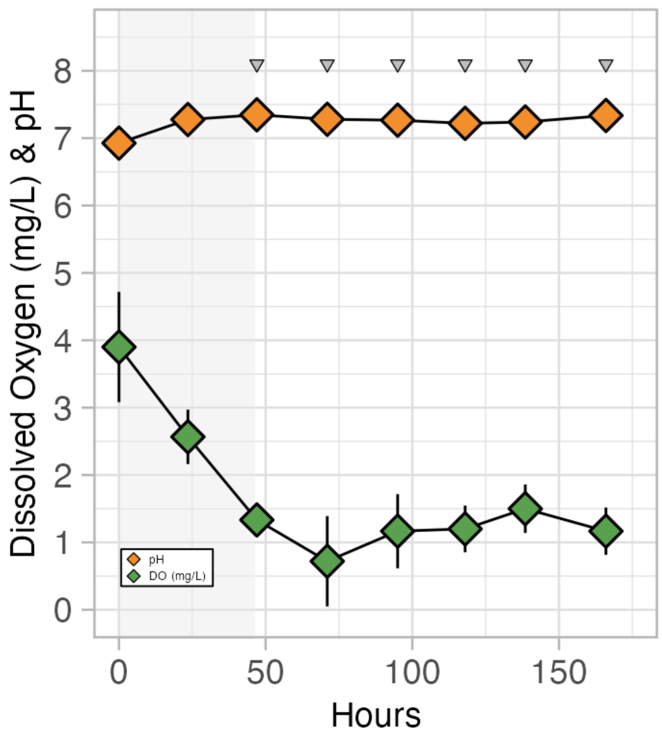


**Supplementary Figure 1. Longitudinal profiles of dissolved oxygen and pH from reactor medium.** Dissolved oxygen and pH (DO [1 mg/L = 1 ppm], green diamonds) from the recirculating media inside triplicate FBRs was measured daily benchtop probes. The grey opaque bars indicate the first 48 hours of batch growth in the FBRs prior to fluidization and the grey triangles represent the timepoints when DNA and RNA were sampled. All error bars indicate standard deviation across 3 replicate FBRs (n = 3).

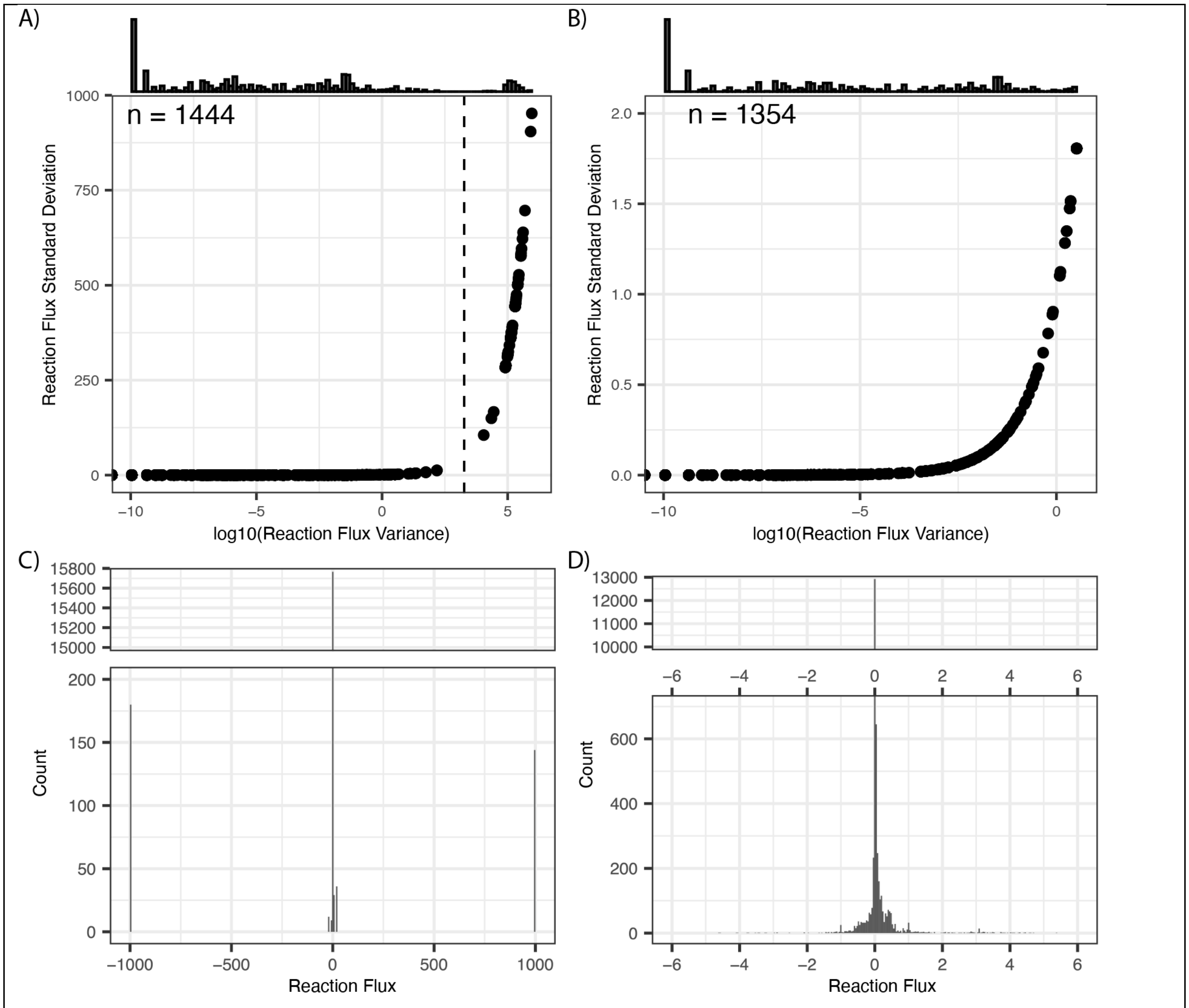


**Supplementary Figure 2. Distribution of FBA reaction fluxes across days and phases and selected cutoffs to reduce complexity and increase confidence. A)** The variance in flux values (mM/gDCW/h) was compared to the standard deviation (Std) for each reaction across each day and phase (n = 12) to identify a confidence threshold (2 Std or 95% confidence intervals, dashed line) for removing highly variable reaction fluxes**. B)** Reactions fluxes below a Std. of 2 that remain after filtering. **C)** Histogram of the non-variable reaction fluxes across all days and phases indicating most reactions fluxes are centered around zero **(D)**.

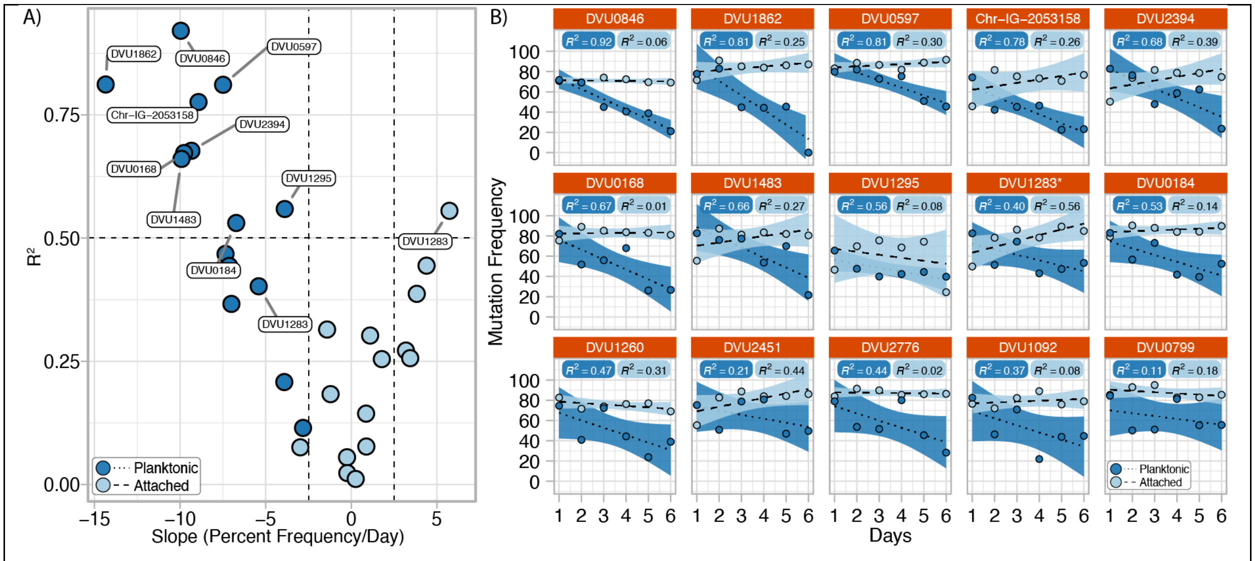


**Supplementary Figure 3. Regression analysis of Dv mutation frequencies over time for each phase. A)** Comparisons of each linear model's coefficient of determination ($R^2$) with its slope for each variant at each phase. Labels were placed on each model that had an $R^2$ greater than .5 and DVU1283 (galU$_{P32S}$). **B)** Each regression model with $R^2$ values for all Dv variants across both phases with ribbons representing the standard error.

**Supplementary Table 1.** Summary of monitored parameters over the experimental time-course

| Monitored Parameters | Description | Instrument/Kit |
|---|---|---|
| Optical Density | OD planktonic Cells | Spectrophotometer |
| Total Nucleic Acids | DNA & RNA | MasterPure Complete DNA & RNA |
| Total Protein | Planktonic and Sediment | B-Per Bacterial Protein Extraction Kit |
| Organics | Lactate, Acetate | IC |
| Headspace gas | Methane | GC (FID/TCD, RGD) |
| pH | | Standard table top probe |
| Dissolved Oxygen | | Standard table top probe |

**Supplementary Table 2.** Workflow for analysis of differential flux states across phases

| Filter Step | Reactions Remaining | Description |
|---|---|---|
| Fluxes for All Common Reactions | 1,444 | Fluxes across all 6 days & 2 phases |
| Reaction Fluxes < 2 Std. | 1356 | Reactions with highly variable fluxes across days and phases have been filtered (n = 88) |
| Reaction Fluxes with no sign flipping | 1346 | FBA fluxes are not on a continuous scale and the sign indicates the direction of the reaction. This step filters reactions that had sign flips and cannot be analyzed as a fold change. (n = 8) |
| Reaction Fold Change (P/S) | 638 | Reactions with non-finite values (Infinite or Nan) generated fold change normalization have been filtered. 705 of these reactions had no flux values, i.e., 0/0 (n = 708) |
| Reaction Fold Change Matrix | 609 | Reactions with no variation, meaning no difference in flux between the two states have been filtered. (n = 29) |
| Reaction Matrix for **Fig. 4D** | 503 | Reactions associated with “Other” & “Biomass” have been filtered. (n = 106) |
